# Auditory brainstem response latencies, but not amplitudes, are associated with gray matter volumes across the human auditory pathway in older adults

**DOI:** 10.64898/2026.08.21.746342

**Authors:** Simón San-Martín, Cristian Aedo, Victor Vidal, Alexis Leiva, Carolina Delgado, Paul H. Délano, Vicente Medel

## Abstract

**Introduction:** Auditory brainstem responses (ABRs) are routinely used to assess neural timing and function along the auditory pathway. In older adults, however, peripheral hearing loss, central auditory dysfunction, and broader structural changes in the brain may converge to shape the recorded response. Because ABR waves arise from multiple overlapping neural sources, how their electrophysiological features map onto specific auditory pathway structures in vivo remains poorly understood. Here, we examined the associations between cortical and subcortical gray matter volumes and the latencies and amplitudes of click evoked ABR Waves I and V in older adults.

**Methods:** We evaluated 88 adults aged > 65 years from the Auditory and Dementia Study (ANDES) cohort. Click evoked ABRs were recorded at 80 dB nHL, and the latencies and amplitudes of Waves I and V were measured. High resolution 3T structural MRI data were processed using voxel based morphometry and standardized anatomical masks to estimate bilateral gray matter volumes of the cochlear nucleus, superior olivary complex, inferior colliculus, medial geniculate nucleus, and auditory cortex. Associations were assessed using partial correlations adjusted for age, pure tone hearing thresholds, and intracranial volume, as well as multivariate linear regression models.

**Results:** ABR latencies, rather than amplitudes, showed significant associations with regional gray matter volumes. After adjustment for age, hearing thresholds, and intracranial volume, larger superior olivary complex volume was associated with shorter Wave I latency (ρ_partial_ = −0.305, p = 0.005), whereas larger medial geniculate nucleus and auditory cortex volumes were associated with shorter Wave V latency (ρ_partial_ = −0.265, p = 0.014 and ρ_partial_ = −0.404, p < 0.001, respectively). In multivariate models, superior olivary complex volume remained associated with Wave I latency (β = −0.310, p = 0.007). Medial geniculate nucleus volume was initially associated with Wave V latency (β = -0.247, p = 0.038); however, this relationship was attenuated once auditory cortex volume was included in the model (β = -0.350, p = 0.002), which emerged as the dominant predictor. Inferior colliculus volume was not significantly associated with Wave V latency or amplitude.

**Conclusions:** In older adults, ABR latencies showed selective associations with regional gray matter volumes, whereas amplitudes did not. These associations extended beyond the structures traditionally considered the main generators of Waves I and V, suggesting that interindividual variation in ABR latency may reflect distributed anatomical variation across the auditory pathway rather than a strict one-wave-one-generator correspondence.

## Introduction

Auditory Brainstem Responses (ABRs) are scalp recorded electrophysiological signals evoked by brief acoustic stimuli. They are commonly described as consisting of five to six vertex positive waves occurring within the first 10 ms after stimulus onset, reflecting the synchronized activation of ascending auditory nerve fibers and brainstem auditory pathway nuclei (Eggermont, 2019; Jewett et al., 1970). Because they are clinically proposed to provide relevant information about neural timing, conduction velocity, synchrony and general pathway function, ABRs have long been routinely used in clinical and research settings to assess auditory function, including sensorineural hearing loss (Simmons & Smith, 1982), auditory neuropathy (Berlin et al., 2010; Starr & Rance, 2015), retrocochlear disorders (Starr et al., 1996), intraoperative monitoring (Bubeníková et al., 2026; Wazen, 1994), and neonatal hearing assessment (Gorga et al., 1989; Thompson et al., 2001). Together, these features have made ABRs a widely used tool for characterizing auditory pathway function across different clinical and physiological conditions.

Aging provides a particularly relevant context for studying ABRs because peripheral auditory decline, central auditory dysfunction, and broader structural changes in the brain occur simultaneously. Cochlear damage and auditory nerve degeneration can alter early components of the response (Turner et al., 2022), whereas reduced neural synchrony and slower conduction across central auditory pathways may affect later waves beyond what can be explained by peripheral hearing thresholds alone (Boettcher, 2002; Konrad-Martin et al., 2012; Pürner et al., 2022). Accordingly, older adults commonly show prolonged ABR latencies and reduced amplitudes (Chu, 1985; Aedo-Sanchez et al., 2023; Maass et al., 2024), but similar electrophysiological changes may arise from different combinations of peripheral and central alterations. This distinction is clinically relevant, as age related hearing loss and speech in noise difficulties are associated with cognitive decline and increased risk of dementia (Anderson et al., 2010, 2012; Delano et al., 2020; Jiang et al., 2022; Kolo et al., 2025, McEvoy et al., 2023). However, the extent to which variability in ABR features in older adults reflects peripheral hearing loss, changes within central auditory structures, or broader age related neurodegeneration remains poorly understood.

The electrophysiological nature of ABRs also limits a strict one-to-one correspondence between individual waves and specific anatomical structures. As scalp recorded far field potentials, ABR waves reflect the summed activity of multiple, partially overlapping neural sources rather than the activity of a single auditory nucleus (Jewett & Williston, 1971; Martin et al., 1995). Evidence from animal and computational models supports this distributed origin, showing that disrupting or modifying a single auditory structure can alter several components of the ABR waveform (Britt & Rossi, 1980; Chen & Chen, 1991; Li et al., 2026; Melcher et al., 1996; Manis & Campagnola, 2018; Verhulst et al., 2015). In humans, intracranial recordings have provided functional information about these contributions, linking Wave I mainly to the distal auditory nerve while showing that later waves arise from overlapping activity across brainstem and midbrain auditory structures (Martin et al., 1995; Møller & Jannetta, 1981; Møller et al., 1988). Consequently, it remains unclear how interindividual variation in the anatomy of the auditory pathway relates to specific electrophysiological properties of scalp-recorded ABR waves in vivo.

Here, we investigated how gray matter volumes across subcortical and cortical structures of the auditory pathway relate to the latencies and amplitudes of click evoked ABR Waves I and V in a cohort of older adults. By combining structural MRI morphometry with scalp recorded electrophysiology, we aimed to characterize whether interindividual variation in these electrophysiological properties is associated with specific auditory structures or with a more distributed anatomical pattern across the auditory pathway.

## Methods

### Participants

A sample of 88 older adults (aged 65 years or older) was recruited from the Auditory and Dementia Study (ANDES) cohort, a prospective longitudinal study based in Santiago, Chile (Medel et al., 2024; San-Martin et al., 2025). All participants underwent comprehensive audiological, neuropsychological, and structural magnetic resonance imaging (MRI) evaluations at the Hospital Clínico de la Universidad de Chile. Inclusion criteria for the ANDES cohort were: (1) age >= 65 years, (2) cognitive and functional performance within normal limits as indicated by a Mini Mental State Examination and Pfeffer Activities Questionnaire (FAQ), (3) exclusion of secondary or non age-related causes of hearing loss, and (4) absence of prior or current hearing aid use. Participants with a history of stroke, brain tumors, severe psychiatric disorders, or major neurological conditions were excluded. All participants provided written informed consent prior to participation, and the study protocol was approved by the Institutional Ethics Committee of the University of Chile (OAIC 752/15).

### Pure-Tone Audiometry

To evaluate hearing thresholds and rule out conductive hearing loss, which is known to prolong ABR wave latencies (Ferguson et al., 1998; McGee & Clemis, 1982), a pure-tone audiometry test was performed using a clinical audiometer (AC40, Interacoustics®) in a soundproof room. Pure-tone air conduction thresholds were determined separately for each ear across frequencies ranging from 0.125 to 8 kHz (0.125, 0.25, 0.5, 1, 2, 3, 4, 6, and 8 kHz). A four frequency pure-tone average (PTA; 0.5, 1, 2, and 4 kHz) was calculated for each ear, and the better ear PTA was retained for subsequent analyses, with higher values reflecting greater hearing impairment.

### Auditory Brainstem Responses (ABR)

Auditory brainstem responses (ABRs) were recorded using the Eclipse platform with the EP25 research module (Interacoustics®, Middelfart, Denmark). Conventional 100 μs clicks were delivered monaurally at a suprathreshold intensity of 80 dB nHL through EARTone 3A insert earphones. Stimuli were presented with alternating polarity at a repetition rate of 21.1 Hz. Recording electrodes were placed at Fz as the active electrode, over the ipsilateral and contralateral mastoid processes (M1 and M2), with the ground electrode positioned over the right supraorbital region. Electrode impedances were maintained below 5 kΩ. Signals were band-pass filtered between 0.15 and 3 kHz, and 2,000 artifact free sweeps were averaged for each recording. Two replicate waveforms were obtained to confirm the reproducibility of the response. Waves I and V were identified from the replicated traces. Latency was defined as the time from stimulus onset to the corresponding positive peak, and amplitude was measured from each positive peak to its subsequent trough, following established protocols for click evoked ABRs in older adults (Delano et al., 2020; Maass et al., 2024). Measurements obtained from the left and right ears were averaged to derive bilateral Wave I and Wave V latency and amplitude values for each participant. When Wave V was present but Wave I was not detectable, its amplitude was assigned a floor value of 0.02 μV, corresponding to the lowest measurable amplitude, following the procedure described by Delano et al., 2020.

### MRI acquisition and analysis

Structural MRI was acquired using a 3-Tesla MAGNETOM Skyra scanner (Siemens Healthcare GmbH, Erlangen, Germany) with a 32-channel head coil. High resolution T1-weighted images were obtained using a three dimensional magnetization prepared rapid gradient echo sequence (MPRAGE; repetition time = 2,300 ms, echo time = 2.3 ms, flip angle = 8°, 176 slices, matrix size = 256 × 256, voxel resolution = 0.94 × 0.94 × 0.9 mm³).

T1 weighted images were processed using FreeSurfer version 6.0 through the standard recon-all pipeline. Cortical parcellation was performed according to the Desikan–Killiany atlas. FreeSurfer was also used to estimate total intracranial volume (eTIV), which was included as a covariate to account for individual differences in head size. All cortical segmentations were visually inspected following standard quality control procedures.

Subcortical auditory volumes were estimated using voxel-based morphometry in SPM12: the Cochlear Nucleus (CN), the Superior Olivary Complex (SOC), the Inferior Colliculus (IC), the Medial Geniculate Body of the Thalamus (MGN), and the Primary Auditory Cortex (AC; corresponding to the bilateral transverse temporal gyrus of Heschl). T1 weighted images were segmented into gray matter, white matter, and cerebrospinal fluid, and the gray matter images were used to create a study specific DARTEL template. Individual maps were normalized to Montreal Neurological Institute space at 1 mm isotropic resolution, modulated to preserve local tissue volume, and smoothed with a 2 mm full-width at half-maximum Gaussian kernel, following our previous VBM procedure (San-Martín et al., 2025; Vidal et al., 2026). Bilateral auditory nuclei were defined using masks from the probabilistic subcortical auditory atlas of Sitek et al. (2019). Regional volumes were calculated by summing voxelwise modulated gray matter values within each mask, after which left and right volumes were averaged for each structure.

### Statistical analysis

Descriptive statistics were calculated for demographic, clinical, electrophysiological, and neuroanatomical variables. Continuous variables are reported as mean ± standard deviation and range, whereas categorical variables are reported as counts and percentages. Variable distributions were evaluated through visual inspection and the Shapiro–Wilk test.

Associations among age, better ear PTA, ABR latencies and amplitudes, and regional gray matter volumes were first examined using Spearman rank correlations. Structure–function relationships were then assessed using partial Spearman correlations between bilateral auditory region volumes and the latencies and amplitudes of Waves I and V, controlling for age, PTA, and estimated total intracranial volume (eTIV).

Post-hoc ordinary least squares regression analyses were conducted for latency measures that showed significant adjusted correlations with regional gray matter volumes. All continuous variables were standardized before model estimation. Wave I latency was modeled as a function of the volume of Superior Olivary Complex, age, PTA, and eTIV. Wave V latency was first modeled as a function of medial geniculate nucleus volume and the same covariates. Bilateral transverse temporal gyrus volume was then added in a second model to examine the relative contributions of subcortical and cortical anatomy. Multicollinearity was assessed using variance inflation factors, with values below 2 considered acceptable, and model assumptions were evaluated through residual diagnostics and inspection of influential observations. Given the exploratory nature of the study, p-values were not adjusted for multiple comparisons and should be interpreted accordingly. All statistical tests were two-tailed, with significance defined as p < 0.05.

## Results

The final sample included 88 older adults, of whom 56 were female (63.6%). Participants had a mean age of 73.5 ± 5.2 years and a mean education of 9.6 ± 4.2 years. Mean MMSE and GDS scores were 28.0 ± 1.3 and 3.1 ± 3.2, respectively. Better ear PTA was 28.8 ± 11.6 dB HL; subjects had normal hearing to moderate hearing loss accordingly. Mean estimated total intracranial volume was 1,359 ± 129 cm³. Full demographic and clinical characteristics are presented in Table 1.

**Table 1:** Demographic and Clinical Characteristics of the Study Cohort (N = 88)

| Variable | Mean / N | SD / % | Median | Minimum | Maximum | Range |
| --- | --- | --- | --- | --- | --- | --- |
| Age (years) | 73.49 | 5.20 | 73 | 65 | 85 | 20 |
| Sex: Female | 56 | 63.64% | - | - | - | - |
| Sex: Male | 32 | 36.36% | - | - | - | - |
| Schooling (years) | 9.61 | 4.22 | 11 | 0 | 20 | 20 |
| Mini Mental State Examination (MMSE) | 27.99 | 1.33 | 28 | 22 | 30 | 8 |
| Geriatric Depression Scale (GDS) | 3.14 | 3.18 | 2 | 0 | 14 | 14 |
| Pure Tone Average (PTA, dB HL) | 28.82 | 11.58 | 27.5 | 7.5 | 53 | 45.5 |
| eTIV (cm<sup>3</sup>) | 1,358.98 | 129.02 | 1,359.03 | 1,082.79 | 1,659.48 | 576.69 |

We first characterized the gray matter volumes of the auditory regions included in the analysis. Among the subcortical regions, the medial geniculate nucleus showed the largest mean bilateral volume (0.2034 ± 0.0310 cm³), followed by the inferior colliculus (0.1896 ± 0.0251 cm³), cochlear nucleus (0.1044 ± 0.0166 cm³), and superior olivary complex (0.0467 ± 0.0102 cm³). The auditory cortex ROI, defined bilaterally as the transverse temporal gyrus, showed a mean gray matter volume of 0.8385 ± 0.1303 cm³. Full volumetric characteristics are presented in Table 2.

**Table 2:** Volumetric Characteristics of Auditory Pathway Structures (N = 88)

| Structure | Mean Volume (cm<sup>3</sup>) | SD (cm<sup>3</sup>) | Median (cm<sup>3</sup>) | Minimum (cm<sup>3</sup>) | Maximum (cm<sup>3</sup>) |
| --- | --- | --- | --- | --- | --- |
| Cochlear Nucleus (CN) | 0.1044 | 0.0166 | 0.1046 | 0.0643 | 0.1429 |
| Superior Olivary Complex (SOC) | 0.0467 | 0.0102 | 0.0455 | 0.0252 | 0.0711 |
| Inferior Colliculus (IC) | 0.1896 | 0.0251 | 0.1861 | 0.1186 | 0.2663 |
| Medial Geniculate Nucleus (MGN) | 0.2034 | 0.0310 | 0.2028 | 0.1180 | 0.2768 |
| Auditory Cortex (AC) | 0.8385 | 0.1303 | 0.8295 | 0.5530 | 1.1355 |

We next examined whether age impacted the auditory pathway. We saw a heterogeneous pattern of age related volumetric decline across the central auditory pathway, where the CN (ρ_partial_ = -0.379, p-value < 0.001), MGN (ρ_partial_ = -0.355, p-value < 0.001), and IC (ρ_partial_ = -0.219, p-value = 0.043) showed negative partial correlations with age after controlling for peripheral hearing thresholds and estimated total intracranial volume. In contrast the SOC (ρ_partial_ = -0.190, p-value = 0.081) and AC (ρ_partial_ = -0.115, p-value = 0.291) did not reveal significant results.

Next we examined whether peripheral hearing sensitivity was related to age, regional anatomy, or ABR properties. As expected, age was positively associated with better ear PTA (ρ_partial_ = 0.429, p < 0.001), indicating poorer hearing thresholds at older ages. Higher PTA (greater hearing loss) was also associated with smaller medial geniculate nucleus volume (ρ_partial_ = −0.233, p = 0.029) and lower Wave I amplitude (ρ_partial_ = −0.241, p = 0.024). No other regional volumes or ABR measures were significantly associated with PTA in the unadjusted analyses. Scatterplots of the associations are shown in the supplementary material Figure S1.

**Figure 1.**
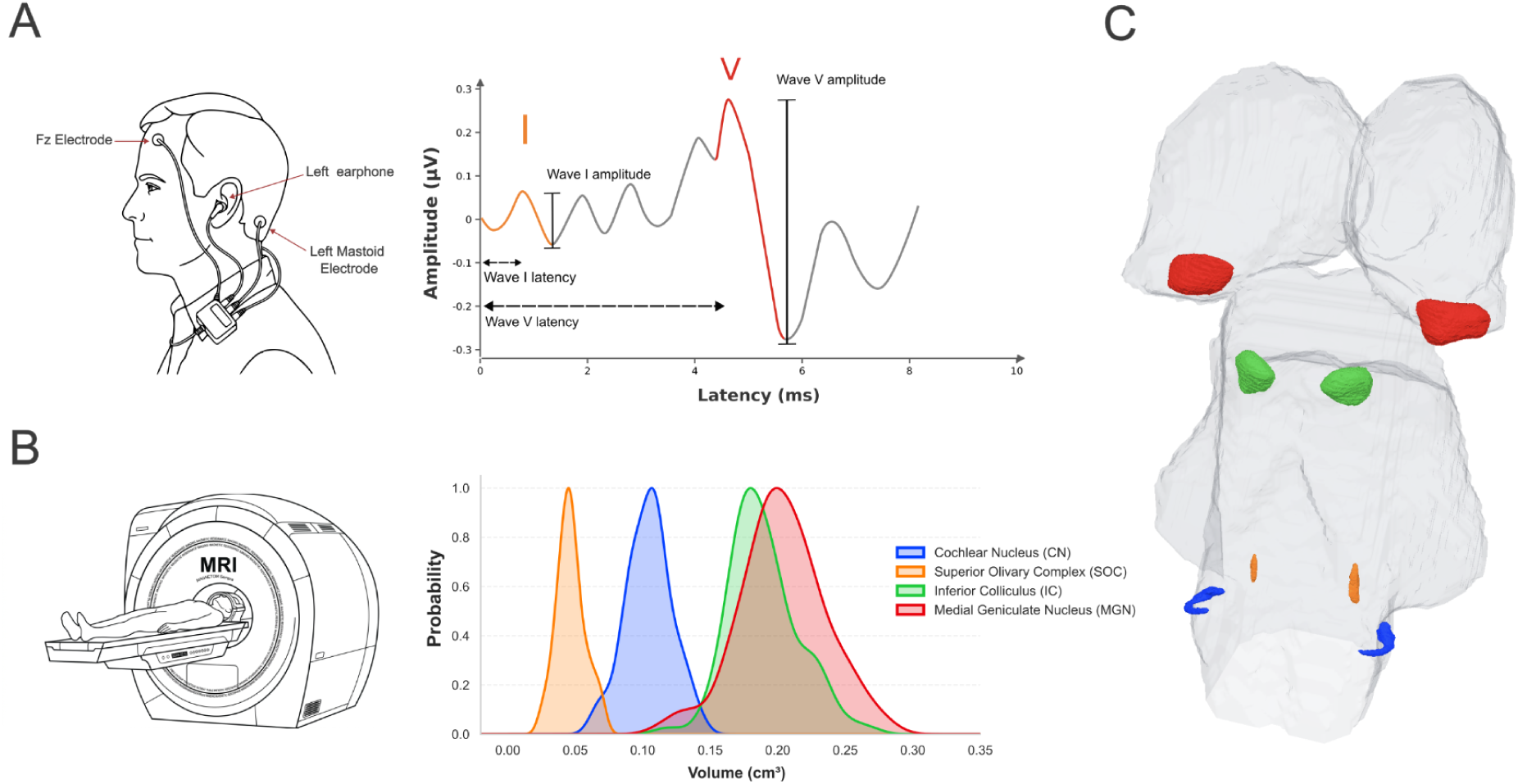
ABR waves of interest and the associated structures in the brainstem for waves I and V found in our analysis. (A) Example of a typical ABR wave showing where latencies and amplitudes are calculated for waves I and V, color indicates associated brainstem structures found in our analysis. (B) Volume distribution of subcortical auditory nuclei (bilateral average) obtained from MRI measurement: Cochlear Nucleus (CN), Superior Olivary Complex (SOC), Inferior Colliculus (IC), and Medial Geniculate Nucleus (MGN). (C) Plot of the average volumes of all subjects for each structure along the brainstem. All volumes are expressed in cubic centimeters (cm³).

Having established these relationships between peripheral hearing sensitivity and age, we next asked whether variation in auditory region volumes was directly related to specific ABR properties.

Interestingly we found that a larger SOC volume was associated with shorter Wave I latency (ρ = -0.30, p-value = 0.005), whereas a larger AC volume was associated with shorter Wave V latency (ρ = -0.23, p-value = 0.027). In contrast, no significant associations were observed between regional gray matter volumes and Wave I or Wave V amplitudes, and no other auditory region volumes were significantly associated with ABR latency measures.

Because age, peripheral hearing sensitivity, and global head size could contribute to these relationships, we next tested whether the observed structure–function pattern remained after adjusting for age, PTA, and eTIV. The association between SOC volume and Wave I latency became stronger after adjustment, with larger SOC volumes associated with shorter Wave I latencies (ρ_partial_ = −0.305, p-value = 0.005). For Wave V, a larger MGN volume was also associated with shorter latency (ρ_partial_ = −0.265, p-value = 0.014). At the same time, the strongest adjusted association was observed for AC volume (ρ_partial_ = −0.404, p-value < 0.001). Importantly, no significant adjusted associations were observed between regional gray matter volumes and Wave I or Wave V amplitudes, reinforcing the selective relationship between auditory region anatomy and ABR timing (Figure 2).

**Figure 2.**
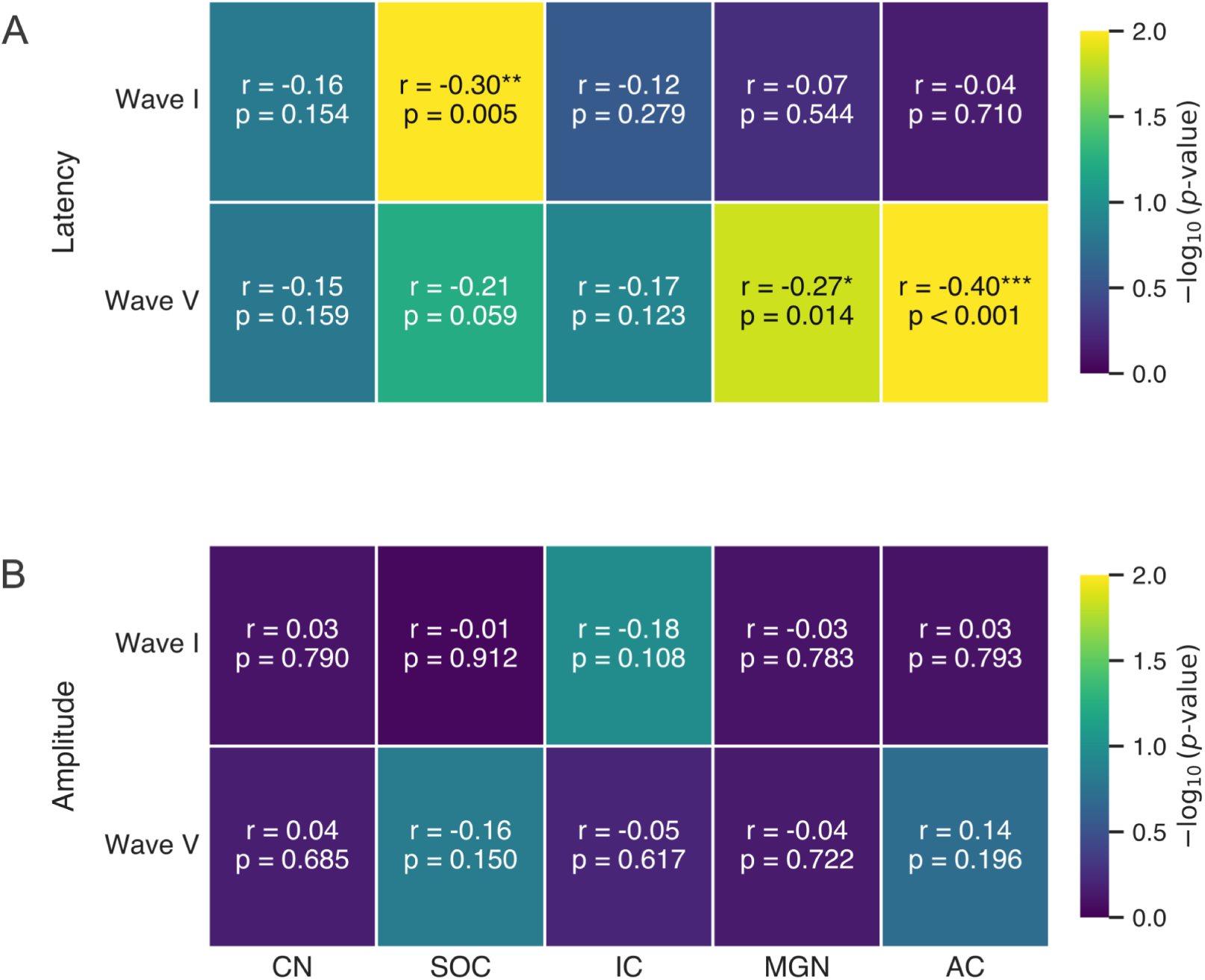
Heatmaps showing partial Spearman correlation (ρ) and p-values (p) for bilateral average ABR latencies (A) and amplitudes (B) for Wave I and Wave V vs. auditory structure volumes (CN, SOC, IC, MGN, AC), controlling for Age, eTIV, and PTA. In the logarithmic color scale, blue colors indicate higher p-value, meanwhile green & yellow colors indicate lower significant p-values. Significance levels: *p < 0.05, **p < 0.01, ***p < 0.001.

To examine whether the adjusted structure and function associations remained after simultaneously accounting by other variables like age, PTA, and eTIV, we conducted post hoc multivariable regression analyses using standardized variables. For Wave I latency, SOC volume remained the only significant anatomical contributor in the model (β = −0.310, 95% CI [−0.533, −0.087], p-value = 0.007). In contrast, age, PTA, and eTIV showed smaller and non-significant contributions (Figure 3A). The model explained 9.9% of the variance in Wave I latency (R² = 0.0993, p-value = 0.066).

**Figure 3.**
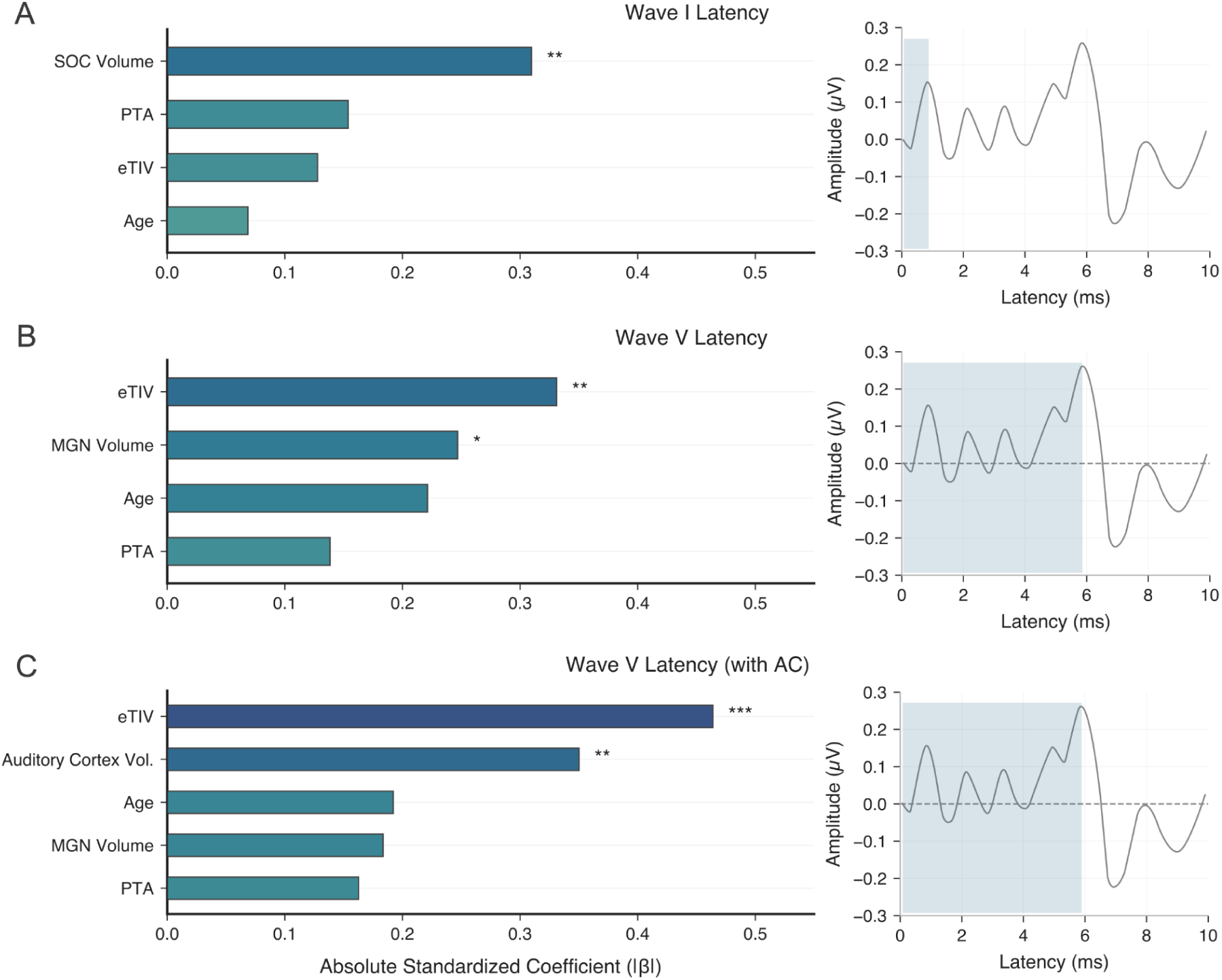
Standardized regression coefficients (β) for OLS regression and the predicted wave latency scheme: (A) predicted Wave I latency based on SOC volume, Age, PTA, and eTIV in the N=88 cohort (R^2^ = 0.099, F = 2.29, p = 0.066). (B) predicted Wave V latency based on MGN volume, Age, PTA, and eTIV (excluding Auditory Cortex) (R^2^ = 0.178, F = 4.48, p = 0.003). (C) predicted Wave V latency based on Auditory Cortex (AC) volume, MGN volume, Age, PTA, and eTIV in the N=88 cohort (R^2^ = 0.269, F = 6.04, p < 0.001). On the right side, the latency of each predicted wave is shown as an example. Significance levels: *p < 0.05, **p < 0.01, ***p < 0.001.

We next examined the relative contributions of subcortical and cortical anatomy to Wave V latency (Figure 3B). In a model including MGN volume, age, PTA, and eTIV, larger MGN volume remained associated with shorter Wave V latency (β = −0.247, 95% CI [−0.480, −0.014], p-value = 0.038). This model explained 17.75% of the variance in Wave V latency (R² = 0.1775, p-value = 0.0025).

When AC volume was added (Figure 3C), AC showed the largest negative anatomical contribution to Wave V latency (β = −0.350, 95% CI [−0.567, −0.133], p-value = 0.002). In contrast, the contribution of MGN volume was attenuated and no longer significant (β = −0.184, p-value = 0.108). The addition of AC volume increased the explained variance from 17.8% to 26.9% (R² = 0.2692, p-value < 0.001). Standardized coefficients for all models are shown in the supplementary Table S2.

## Discussion

In this study, we examined how gray matter volumes across the auditory pathway relate to suprathreshold ABR properties in older adults. The main finding was a selective association with response timing: regional volumes were related to Wave I and Wave V latencies, whereas no significant associations were observed for amplitudes. These relationships remained after accounting for age, peripheral hearing thresholds, and intracranial volume. In the post-hoc regression models, SOC volume retained an independent contribution to Wave I latency, whereas MGN and AC volumes showed large anatomical contributions to Wave V latency. Importantly, this pattern did not follow a strict one wave & one generator correspondence. Furthermore, our results are consistent with previous descriptions indicating that scalp recorded ABRs are composite potentials (Moore, 1987), arising from multiple overlapping sources as demonstrated by both animal studies (Britt & Rossi, 1980; Chen & Chen, 1991; Martin et al., 1995; Melcher et al., 1996, Wang et al., 2025) and human intracranial recordings (Møller & Jannetta, 1981; Møller et al., 1988).

The association between larger SOC volume and shorter Wave I latency is particularly notable because Wave I is commonly attributed to activity in the distal auditory nerve (Møller & Jannetta, 1981; Caird & Klinke, 1987; Biacabe et al., 2001; Jacxsens et al., 2022). Although the overall model did not reach statistical significance (Figure 3), it trended toward significance. Furthermore, the robust partial correlations suggest a meaningful association between these two measures. One possible interpretation involves the medial olivocochlear system (Elgueda & Delano, 2020; Robles & Delano, 2008). Medial olivocochlear neurons originate in the SOC and project to the cochlea, where they modulate outer hair cell activity and cochlear gain (Delano et al., 2020). Through this pathway, variation in SOC anatomy could be related to the timing and synchronization of auditory nerve activity. However, the present study did not include a direct measure of olivocochlear function. The observed relationship should therefore be considered anatomically compatible with an efferent contribution, rather than evidence that SOC volume directly determines Wave I latency.

Wave V showed a different and more distributed anatomical pattern. Its latency was associated with both MGN and AC volumes, but not with IC volume, despite the IC being commonly considered a major contributor to Wave V (Biacabe et al., 2001; Caird & Klinke, 1987; Jacxsens et al., 2022). This could be in part related to smaller variability (or reduced plastic changes) across subjects at the IC specific level. Indeed, previous research demonstrates that age related structural changes along the auditory pathway are more prominent in cortical regions, such as Heschl’s gyrus and the planum temporale, than in subcortical structures (Profant et al., 2014). Nevertheless, one recent study showed a clear reduction in volumetric measures at the IC related to aging variables (Kwok et al., 2025). Moreover, estimated total intracranial volume (eTIV) emerged as the strongest positive predictor of Wave V latency (β = 0.464, p < 0.001). This positive association reflects a straightforward physical principle: a larger intracranial volume corresponds to greater head size and longer physical axonal distances along the brainstem and thalamocortical pathways, naturally resulting in longer conduction times to reach far field scalp electrodes.

One possible interpretation of this departure from the classical generator based account (Melcher et al., 1996; Moore, 1987) is that ABR latencies may reflect not only the properties of their putative local generators, but also structural relationships with subsequent stages of the ascending auditory hierarchy (Biacabe et al., 2001). Under this target centric interpretation, Wave I latency could reflect transmission efficiency and efferent regulation involving its downstream brainstem target, the SOC. In contrast, Wave V latency could reflect cumulative transmission and corticofugal calibration across later thalamocortical targets, including the MGN and AC. Furthermore, because far field ABRs reflect the algebraic summation of overlapping neural sources, Wave V latency may capture cumulative variation across the central auditory pathway rather than the local gray matter volume of the IC alone (Jewett & Williston, 1971; Melcher et al., 1996). Consistent with this interpretation, AC volume showed the largest negative anatomical contribution in the final regression model, while the contribution of MGN volume was attenuated after AC was included. Corticothalamic and corticofugal projections provide a plausible anatomical framework for understanding these associations (Caspary et al., 2008; Da Costa et al., 2021; Sitek et al., 2019), but the present findings cannot determine whether cortical anatomy directly modulates brainstem timing or instead reflects broader variation in central auditory or global brain integrity.

Some limitations should be considered. First, the cross sectional design does not allow us to establish the direction or mechanism of the observed associations. Second, regional gray matter volume is an indirect and relatively nonspecific measure of anatomical integrity, particularly for small subcortical nuclei, as estimated from conventional 3T MRI. Third, we selected the primary auditory cortex as the sole cortical source for measurement, which may introduce a potential bias. Fourth, averaging bilateral volumes and ABR measures across hemispheres and ears may obscure lateralized relationships within the auditory pathway. Finally, we did not include direct measures of white matter integrity, olivocochlear function, or corticofugal connectivity. Despite these limitations, our results show that ABR latencies, rather than amplitudes, are selectively associated with gray matter variation across cortical and subcortical auditory regions in older adults. Given that ABR amplitudes can be particularly sensitive to skull thickness, electrode placement, and physiological noise (Prasher & Gibson, 1980, Møller & Jannetta, 1981; Møller et al., 1988), latencies may provide a more stable electrophysiological correlate of the distributed auditory system’s anatomy.

## Acknowledgment

We acknowledge the support of the ANDES research team involved in data acquisition. V.M. is supported by ANID/FONDECYT de Iniciación 11251578, ANID/FONDECYT Exploración 13240170, and AARG-25-1490306 from the Alzheimer’s Association. CD is funded by ANID grants ID25I10379 and ID2110096. PHD is funded by ANID grants AC3E CIA250006, ANID/FONDECYT Regular 1261514, FONDEF grant ID25I10328, and Fundación Guillermo Puelma.

## Supplementary Material

**Figure S1:**
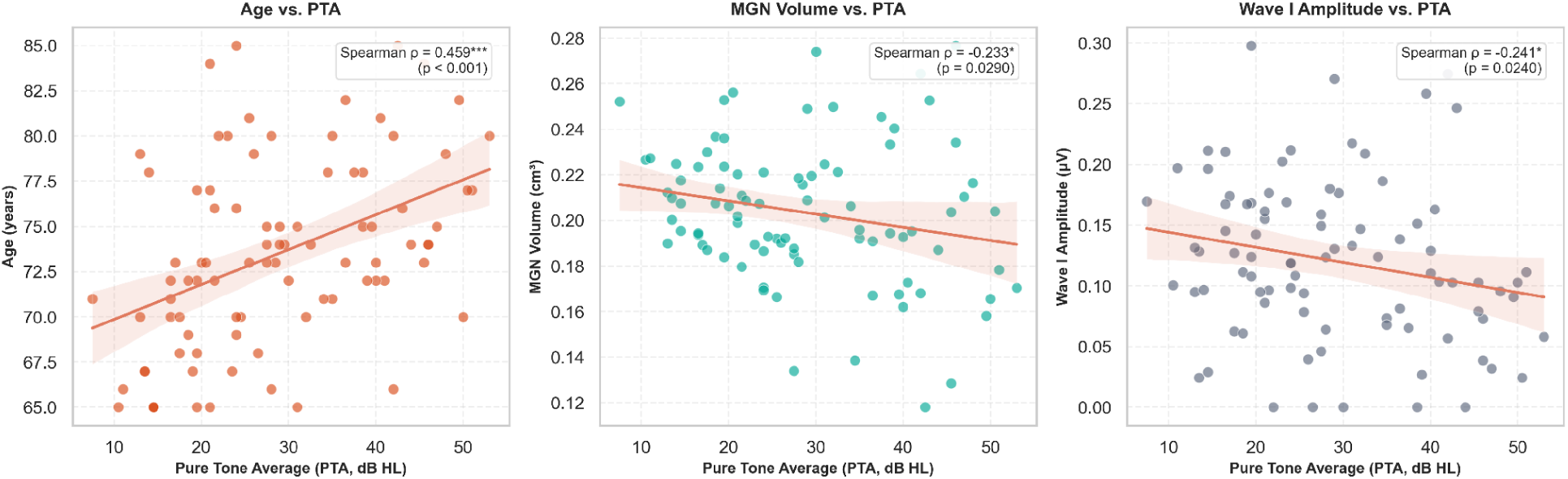
Bivariate Spearman scatterplots and linear trend lines for significant associations with Pure Tone Average (PTA) in the N=88 cohort: Age vs. PTA, Medial Geniculate Nucleus (MGN) volume vs. PTA, and Wave I amplitude vs. PTA.

**Table S1:**
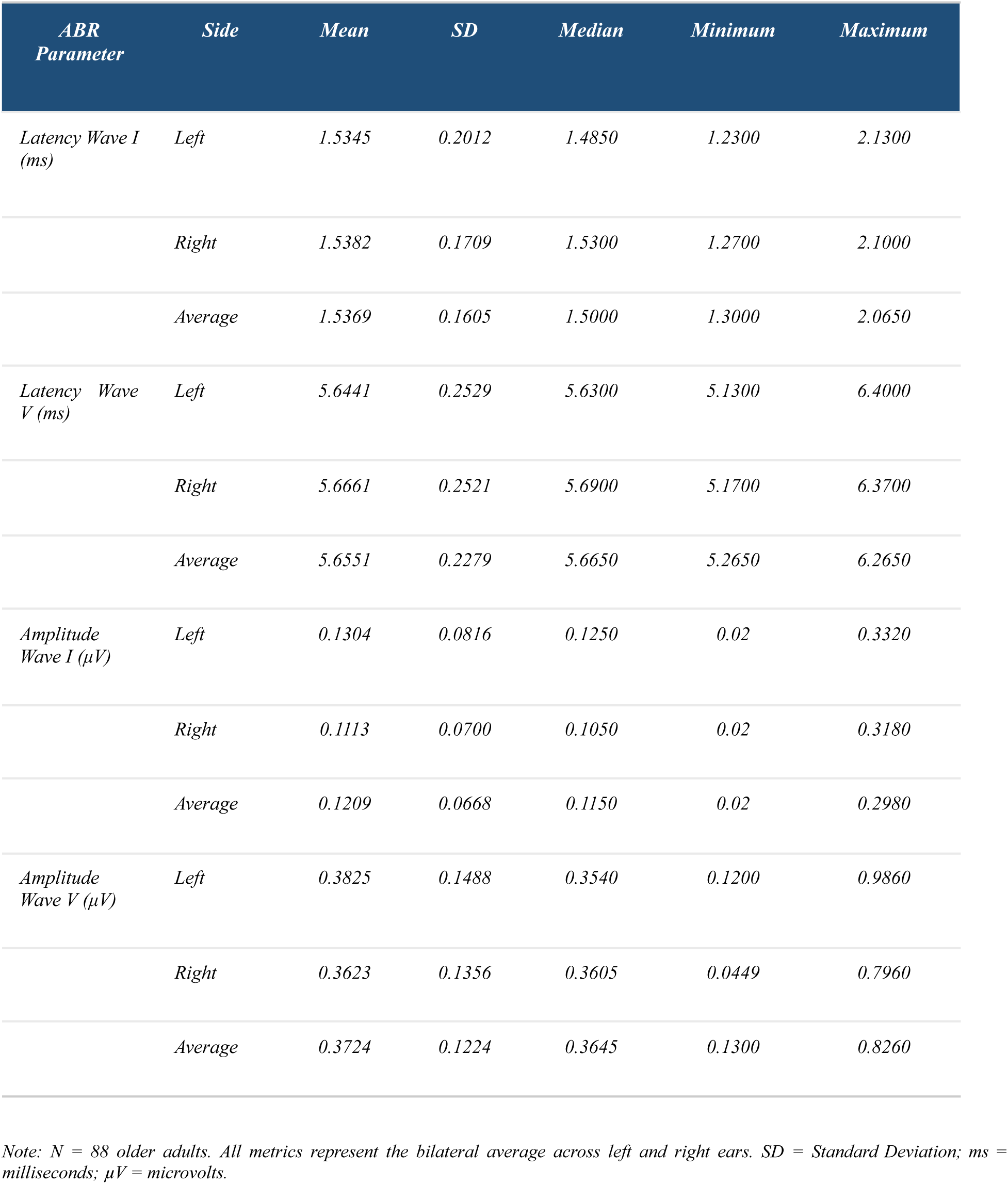
Electrophysiological Basal ABR Parameters (N = 88)

**Table S2:**
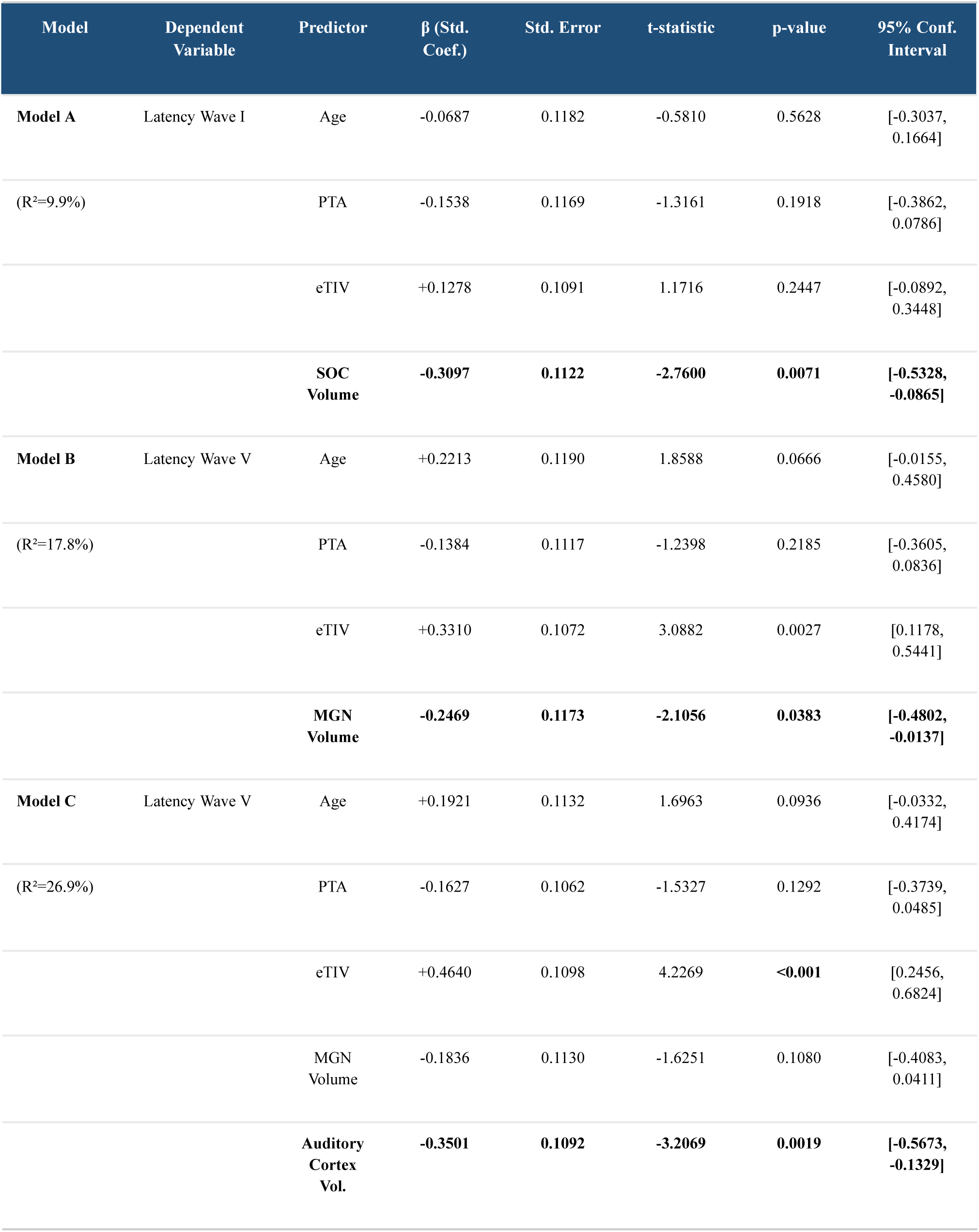
Summary of Multivariate OLS Regression Models (N = 88)

